# Label-Free Melanoma Phenotype Classification Using Artificial Intelligence-Based Morphological Profiling

**DOI:** 10.1101/2024.06.28.601235

**Authors:** Evelyn Lattmann, Andreja Jovic, Julie Kim, Tiffine Pham, Christian Corona, Zhouyang Lian, Kiran Saini, Manisha Ray, Vivian Lu, Aizhan Tastanova, Stephane C. Boutet, Mitchell P. Levesque

**Affiliations:** University of Zurich, University Hospital of Zurich, Department of Dermatology, Schlieren, Switzerland; Deepcell Inc., Menlo Park, CA

## Abstract

Melanomas are the deadliest skin cancers, in part due to cellular plasticity and heterogeneity. Intratumoral heterogeneity drives varied mutable phenotypes, specifically “melanocytic” and “mesenchymal” cell states, which result in differential functional properties and drug responses. Definitive and rigorous classification of these phenotypic states has been challenging with conventional biomarker-based methods, and high-parameter molecular methods are cell-destructive, labor-intensive, and time-consuming. To overcome these technical and practical limitations, we utilized label-free artificial intelligence-based morphological profiling to classify live melanoma cells into melanocytic and mesenchymal phenotypes based on high resolution imaging of single cells.

To predict the phenotypes of single melanoma cells based on morphology alone, we developed the AI-based ‘Melanoma Phenotype Classifier’ trained with 19 patient-derived cell lines with known melanocytic or mesenchymal transcriptional profiles. To link phenotypic state with high-dimensional morphological profiles, cells were subjected to genetic and chemical perturbations known to shift phenotypic states. The AI classifier successfully predicted phenotypic shifts which were confirmed by single-cell RNA-Seq (scRNA-Seq). These results demonstrate that correlations between melanoma cell phenotypes and morphological changes are detectable by AI. Additionally, the Melanoma Phenotype Classifier was applied to dissociated tumor biopsy samples and characterization of phenotypic heterogeneity was supported by scRNA-Seq transcriptional profiles.

This work establishes a link between cell morphology and melanoma phenotypes, laying the groundwork for the use of a label-free morphology-based method for phenotyping live melanoma cells combined with additional analyses.

## Introduction

Cellular phenotyping offers crucial insights into the relationships between interacting molecular networks and cellular functions. In cancer, phenotyping is important in characterizing cellular heterogeneity. Melanoma, known for its aggressive nature and propensity for rapid progression, exemplifies the urgent need for innovative phenotyping methods. The plasticity of melanoma cells enables a range of cellular transitions leading to cellular heterogeneity, metastasis, and resistance to therapy. Melanoma cells can transition from a differentiated melanocytic phenotype through intermediate states to a less differentiated mesenchymal phenotype (1–4). Phenotype switching is governed by the microphthalmia-associated transcription factor (MITF), a key regulator of pigmentation (1,5–7). High MITF levels are associated with a melanocytic phenotype, whereas reduced MITF levels, often triggered by external stimuli such as TGFβ, mark a shift towards a mesenchymal state with an upregulation of pro-invasive genes (8–12). The dedifferentiated, slow-cycling state is the main driver of resistance to both targeted therapies and immune checkpoint inhibitors (13–17). However, upregulation of MITF leading to a phenotypic shift towards a melanocytic phenotype has also been observed as a resistance mechanism, highlighting the critical role of phenotype switching in drug resistance (14,18,19).

Currently, a variety of methods are employed to characterize cellular phenotypes, including molecular techniques (e.g., RNA sequencing), cell biological methods (e.g., flow cytometry, Cytometry Time-Of-Flight), biochemical approaches (e.g., proteomics), and imaging techniques (e.g., high-content screening or cell painting). Despite the information these methods provide in cellular characterization, they are time-consuming, labor-intensive, and cost-prohibitive. Most of these methods also require cell lysis or fixation, which prevent further functional analyses. Moreover, marker gene sets for phenotype assignment are not consistent across laboratories (1,3,20,21). This underscores the necessity for orthogonal approaches that are time-efficient, cost-effective, and enable label-free phenotyping of individual cells.

Given the potential visual differences between melanoma phenotypes (22,23), we hypothesized that morphological analysis could serve as a label-free method for classifying melanoma cell phenotypes while maintaining cellular integrity. For this purpose, we employed Deepcell technology, which combines microfluidics, high resolution brightfield cell imaging, and machine learning (24,25).

The goal of this study was to use high resolution cell imaging and AI to develop a Melanoma Phenotype Classifier to enable label-free phenotyping of melanoma cells at the single-cell level. We used this classifier to distinguish melanocytic from mesenchymal cell states and used transcriptional signatures (scRNA-seq) to measure accuracy of morphology-based predictions, and further tested the classifier in identifying phenotypic shifts induced by chemical or genetic perturbations (Fig. 1). Correlating cellular morphology with these genetic and pharmacological perturbations facilitates a deeper understanding of their impact on cellular responses that can be used for finding targeted treatments. Importantly, we also illustrated its applicability in assessing the phenotypes of clinical samples from melanoma patients. Our findings show that morphology accurately captures complex phenotype information. We foresee that morphotyping (i.e. phenotyping based on morphology) will enable label-free single-cell phenotype-based classification and ultimately sorting of live melanoma cells. Within the broader context of oncology, our approach paves the way for morphotyping-based classification and sorting of a wide range of tumor cells, beyond melanoma, and thus presents a cost-effective alternative to traditional transcriptomic methods.

**Figure 1:**
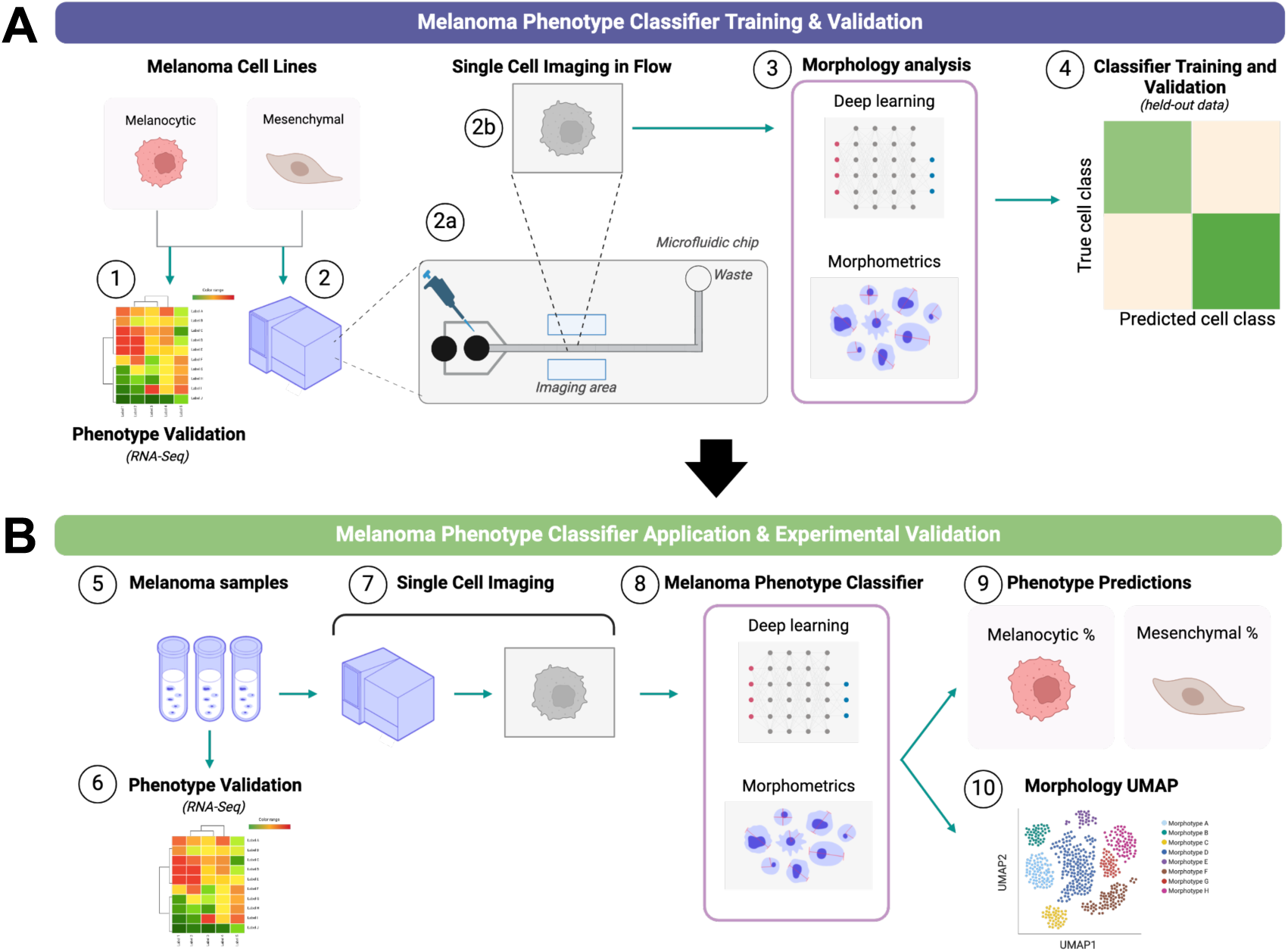
Development of the Melanoma Phenotype Classifier. **(A)** Melanoma cell lines with known melanocytic or mesenchymal phenotypes based on gene expression signatures (*1*) were chosen for morphology analysis on the Deepcell platform (*2*). For each cell line, a single cell suspension was flowed through a microfluidic chip (*2a*). Each cell was imaged using a high-speed, high-resolution camera as it flowed through the imaging area (*2b*). Multi-parameter morphology analysis was performed on each cell image, analyzing both deep learning and computer vision-based morphometric features (*3*). These features were used to train a random forest classifier (the “Melanoma Phenotype Classifier”) to predict cell state. The Melanoma Phenotype Classifier was validated using a separate set of images independent from the training image set (*4*). **(B)** The Melanoma Phenotype Classifier was applied to patient-derived cell lines and melanoma dissociated tumor cells (DTCs) (*5*). Each sample was used for both gene expression analysis (*6*) and Deepcell image analysis (*7*). The same set of features were measured for each cell (*8*), then the Classifier was used to predict the phenotype of each cell (*9*). Features were used to generate a 2D UMAP to visualize subsets of morphotypes (*10*).

## Materials and Methods

### Commercial and primary human melanoma cell lines

Commercial cell lines were either purchased from commercial vendors or obtained from collaborators as indicated in Supplementary Table 1. Primary human cell lines were acquired from the URPP biobank at the University of Zurich from consenting patients of Caucasian origin, with patient ages between 29 and 84 years, and including 15 male patients and 13 female patients. Detailed information of the primary human cell lines is indicated in Supplementary Table 2. The isolation procedure has been previously described (26).

### Dissociated tumor cell samples (DTCs)

All patient biopsies were purchased from Discovery Life Sciences: all patient biopsies were of Caucasian origin, with patient ages ranging from 42 to 87 years, and 4 female patients and 2 male patients. Detailed information of DTCs are indicated in Supplementary Table 3.

### Sample Preparation and imaging on the Deepcell instrument

Cryopreserved samples were thawed in a water bath at 37°C until a small amount of ice was visible in the vial. Cells were then transferred to a 50 mL conical Falcon tube (Corning REF 430829), and 1 mL of RPMI+GlutaMAX (Gibco REF 61870-036) + 10% FBS (Peak Serum Cataolog No. 04A1232) was slowly added over the course of 1 minute to neutralize the freezing medium. An additional 8 mL of medium was added over the next minute for a total volume of 10 mL. The cell mixture was spun down at 300 g for 5 minutes, and the cell pellet was resuspended in the Deepcell sample buffer to reach a concentration of 1e6 cells/mL. The resulting sample was imaged at ∼30 cells/s on the Deepcell instrument. For each sample, tens of thousands of images were collected on one or more Deepcell instruments in duplicate.

### Perturbation experiments

Two types of perturbation experiments were carried out, chemical and genetic. A375 and SkMel28 cells were treated with 5 µM Vemurafenib (Selleck Chemicals, Catalog No.S1267) in full cell medium for 72 h before they were trypsinized, cryopreserved (in liquid nitrogen for at least 24 h), and then imaged on the Deepcell platform; as controls for these perturbations, we used A375 and SkMel28 cells treated with DMSO in full cell medium for 72 h. The second chemical perturbation used was TGF-β (Bio-techne R&D Systems,, Catalog No. 7754-BH-005), which was reconstituted in HCl per the manufacturer’s instructions. SkMel28 cells were treated with 50 ng/mL TGF-β in full cell medium for 72 h before they were trypsinized, cryopreserved (in liquid nitrogen for at least 24 h), and then imaged on the Deepcell platform; we used SkMel28 cells treated with HCl in full medium for 72 h as the control. The cell line M990922 was genetically perturbed by CRISPR knockout of the STK11 gene, as described in Dzung et al (27), and imaged on the Deepcell instrument. As a control, we imaged a clone of M990922 that harbored a non-targeting scrambled (SCR) short-guide RNA construct with wild-type STK11.

### Bulk RNA-Seq

The Melanoma phenotypes were verified by bulk RNA-Seq using the approach originally published in Widmer et al. (28), and recently applied in Eichhoff et al., 2023 (26). The counts are normalized across samples to have a total depth of one million. Using a set of samples with known phenotype as a reference, the expression level from melanocytic markers from each sample was correlated to the averaged melanocytic marker gene expressions from melanocytic reference samples. The correlation analysis was repeated using known mesenchymal samples and mesenchymal marker gene expressions. The Pearson correlation scores to melanocytic expression and mesenchymal expression in test samples were then visualized using a scatter plot, where the distinct clusters of melanocytic and mesenchymal phenotypes were identified. Using a correlation score cutoff value of 0.6, each test sample was phenotyped as either mesenchymal, melanocytic, or intermediate.

### Single cell RNA-Seq

In preparation for scRNA-Seq, cells were counted and resuspended at a concentration of 1000 cells/μL in PBS supplemented with 0.04% BSA. Cells were run using the 10x Chromium Next GEM Single Cell 3’ Reagent Kits v3.1 Dual Index (PN-1000268, 10x Genomics, Pleasanton, CA, United States) per manufacturer guidelines, and all runs were performed in duplicate. 5000 cells were targeted for each library. The cells were then loaded into individual channels on the Chromium Next Gem Chip G (PN-1000127) and encapsulated into gel beads in emulsion (GEMs) via the 10x Chromium iX instrument (PN-1000328). Single cell GEMs were lysed and underwent reverse transcription, resulting in barcoded cDNA. The resulting barcoded libraries underwent amplification, fragmentation, end-repair, adaptor ligation, and sample index attachment using Dual Index Kit TT Set A (PN-1000215). Final single cell transcriptome libraries were sequenced at a minimum depth of 20,000 reads per cell on the Illumina Nextseq 2000 and Novaseq600 instruments as follows: 28 cycles (Read 1), 10 cycles (i7 Index), 10 cycles (i5 index), 90 cycles (Read 2). The sequenced reads were processed using the Cell Ranger analysis pipeline v.7.0.1 (29) to generate FASTQ files and align sequencing reads to a pre-built human reference transcriptome version GRCh38-3.0.0 by 10x genomics. The raw binary format output with cell barcodes and unique molecular identifier count results were used for the downstream analysis.

### Statistical Methods and Hypothesis Testing for single cell RNA-Seq data

Using the Scanpy tookit (30), the binary data in hierarchical data format (H5) from each sample were combined into an AnnData object. Basic filtering was performed based on the number of genes expressed in the count matrix, total counts per cell, and the percentage of counts in mitochondrial genes to reduce the number of poor-quality cells present in the data. Highly variable genes were identified using the built-in function from Scanpy. Doublets were identified using Scrublet (31) and to make each cell comparable, the data was normalized to 10,000 reads per cell and corrected by regressing out using total counts, percentage of the mitochondrial genes expressed and the cell cycle scores. Lastly, the cells expressing high levels of immune markers (e.g. CD45) were excluded from the downstream analysis. For embeddings analysis, the data is corrected using Harmony algorithm (32) to reduce the effects of the sample specific effects, and then the mesenchymal and melanocytic marker expressions data is projected in UMAP. To classify a sample into mesenchymal or melanocytic type based on marker gene expression, the marker genes were selected from Hoek et al (33) and Lischetti et al. (21). For each cell, the scores for mesenchymal and melanocytic phenotypes were calculated using the score genes method available from Scanpy (34), where the distribution of the scores from known melanocytic and mesenchymal samples were used to determine the thresholds categorizing a given cell as either melanocytic or mesenchymal. Each cell is categorized as one of the following: expressing melanocytic marker genes, expressing mesenchymal marker genes, expressing both melanocytic and mesenchymal marker genes, or expressing neither mesenchymal nor melanocytic. Once the phenotype was assigned for each cell, the composition of the assigned phenotypes was calculated for each sample.

### Foundation Model Development

We implemented a variant of the VICReg model to construct the self-supervised deep learning embedding model. In addition, a computer vision algorithm was used to quantitatively extract morphometric features such as cell shape and size. The Deepcell Human Foundation Model (HFM) is a hybrid architecture combining self-supervised learning and computer vision morphometrics. At the time of this study, the HFM training dataset included polystyrene beads, human cancer cell lines, various immune cells, stromal cells, healthy and diseased tissue samples, and iPSC-derived differentiated cells.

### Imaging classification analysis

The images from melanocytic and mesenchymal phenotyped melanoma samples were labeled as melanocytic or mesenchymal respectively, whereas the images from melanoma samples with unknown phenotypes were annotated as unknown. Each image was encoded as a combination of morphometrics and deep learning-based features using the Deepcell Human Foundation Model (HFM). After removing out-of-focus cells and images of debris, images from phenotyped samples were randomly split into training and validation sets using a 4-fold cross-validation method for development and evaluation of the Melanoma phenotype classifier, which was based on a random forest classifier framework. For each cell line from the validation set, the percentage of cells classified as mesenchymal and melanocytic was generated. The melanoma samples in the test sets, coming from the perturbation and DTC experiments, were classified as either mesenchymal or melanocytic phenotype, using the proportion of cell images classified as mesenchymal or melanocytic by the majority voting method.

### Ethics Statement

All experimental protocols in this study involving human subjects were reviewed and approved by the local ethics committee (BASEC 2014-0425), and were conducted in accordance with the Declaration of Helsinki. Informed consent was obtained from all individual participants included in the study.

## Results

### Development and Validation of a Melanoma Phenotype Classifier

We developed an AI-based classifier to identify melanoma cells exhibiting either a melanocytic or mesenchymal phenotype based on their morphologies. Our workflow for training and validating the Melanoma Phenotype Classifier entailed individually imaging cell suspensions of 12 melanocytic and 7 mesenchymal cell lines on the Deepcell platform (Fig. 1A). For each line, phenotypes were verified by bulk RNA-Seq analysis using established melanocytic and mesenchymal transcriptional signatures (21,35). Cell images from the training set were used to generate the classifier independent from the validation set used to evaluate the classifier. Using 4-fold cross-validation, imaging data for the 19 cell lines was divided such that no cell line was part of both training and validation. Images from the training and validation sets were processed by the Deepcell Human Foundation Model (HFM) framework, which quantifies visual attributes of individual cell images through extraction of both morphometric and deep learning features (51 and 64 features, respectively, Supplementary Table 5). The Melanoma Phenotype Classifier predicted the frequency of mesenchymal and melanocytic cells in each sample from the validation set, which harbored differing driver mutations from a wide diversity of tumor sites (Table 1, see also Materials and Methods, Supplementary Table 1, 2). The Classifier predicted phenotype frequencies in high concordance with classification as determined by bulk RNA-Seq (Table 1) and selected samples analyzed with scRNA-Seq (Supplementary Table 6). Of the 12 melanocytic cell lines, all cell phenotype predictions aligned with bulk RNA-Seq classifications, with a range of 60% to 97%. Similarly, of the 7 mesenchymal cell lines, classifier predictions aligned with RNA-Seq, with a range of 66% to 94%. These results demonstrate that morphology reliably differentiates between melanoma phenotypic states in vitro.

**Table 1.**
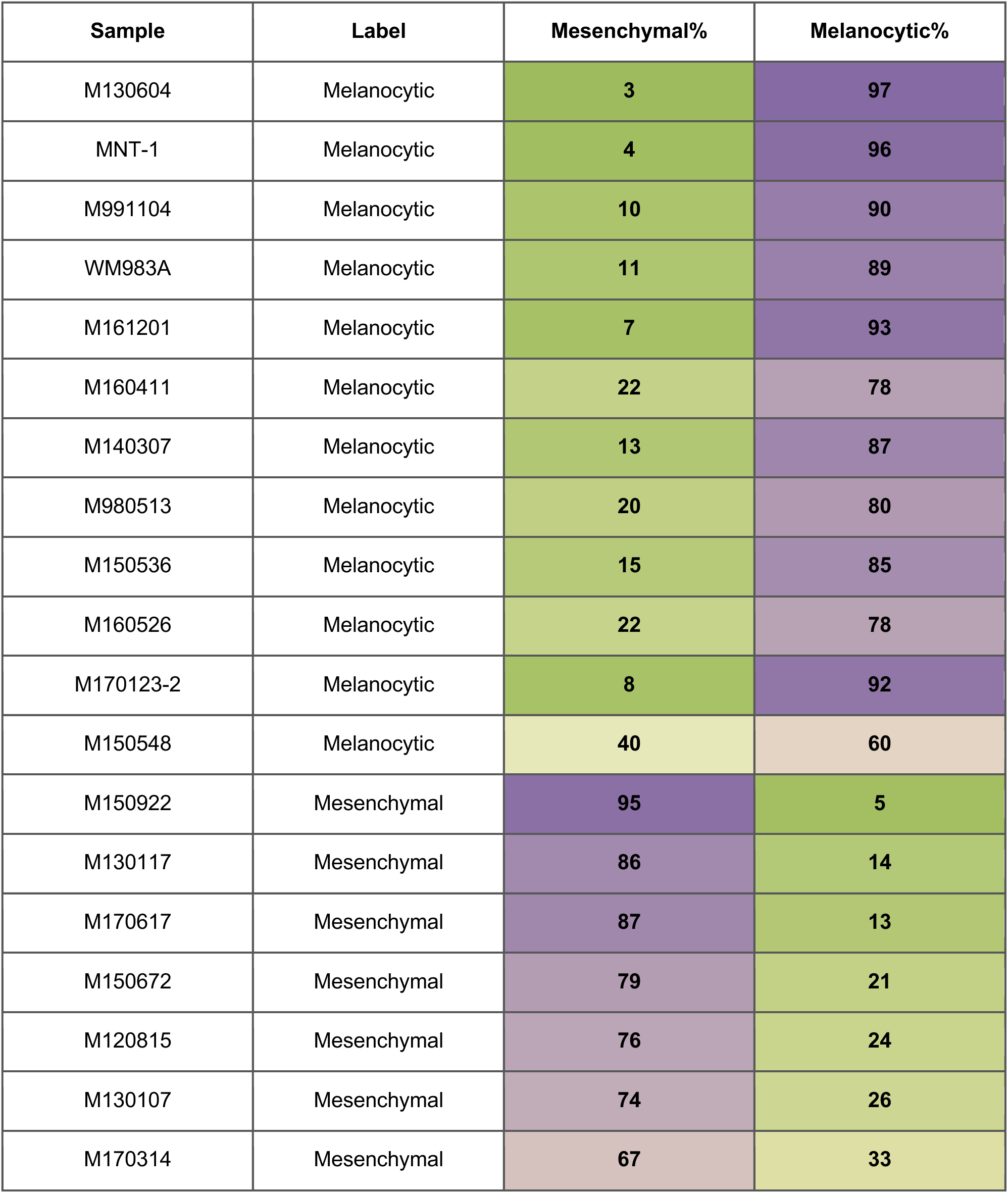
Performance of the Melanoma Phenotype Classifier on the cell lines used for training and validation.

| Sample | Label | Mesenchymal% | Melanocytic% |
| --- | --- | --- | --- |
| M130604 | Melanocytic | 3 | 97 |
| MNT-1 | Melanocytic | 4 | 96 |
| M991104 | Melanocytic | 10 | 90 |
| WM983A | Melanocytic | 11 | 89 |
| M161201 | Melanocytic | 7 | 93 |
| M160411 | Melanocytic | 22 | 78 |
| M140307 | Melanocytic | 13 | 87 |
| M980513 | Melanocytic | 20 | 80 |
| M150536 | Melanocytic | 15 | 85 |
| M160526 | Melanocytic | 22 | 78 |
| M170123-2 | Melanocytic | 8 | 92 |
| M150548 | Melanocytic | 40 | 60 |
| M150922 | Mesenchymal | 95 | 5 |
| M130117 | Mesenchymal | 86 | 14 |
| M170617 | Mesenchymal | 87 | 13 |
| M150672 | Mesenchymal | 79 | 21 |
| M120815 | Mesenchymal | 76 | 24 |
| M130107 | Mesenchymal | 74 | 26 |
| M170314 | Mesenchymal | 67 | 33 |

We next sought to assess morphological diversity of the cell lines with quantitative characterization of morphological differences between the two phenotypes (Fig. 1B, Supplementary Fig. 1). For each cell line, “embeddings” were created using the quantitative morphometric and deep learning features extracted by HFM, effectively creating a representation of each individual cell image. These embeddings were visualized with uniform manifold approximation and projection (UMAP) (36), providing a 2D representation of the “morphology space” for the 19 cell lines whose morphological diversity was highlighted by the use of Leiden clustering (37) (Fig. 2A). The phenotype density representation of the same UMAP (Fig. 2B, Supplementary Fig. 1-2) revealed that each melanoma phenotype occupied a unique part of morphological space, underscoring the morphological differences between the two. Interestingly, cells with intermediate melanoma phenotypes occupied morphology space in the UMAP between that of melanocytic and mesenchymal phenotypes (Supplementary Fig. 2). Representative images of the latter phenotypes from several cell lines further demonstrated the existence of morphological differences (Fig. 2B), including the presence of a subpopulation of pigmented cells in cell line M991104 (Fig. 2B, red circle). As a reflection of these morphological differences in phenotype, the Melanoma Phenotype Classifier exhibited an overall accuracy >80% across the phenotyped 19 cell lines (Fig. 2C).

**Figure 2:**
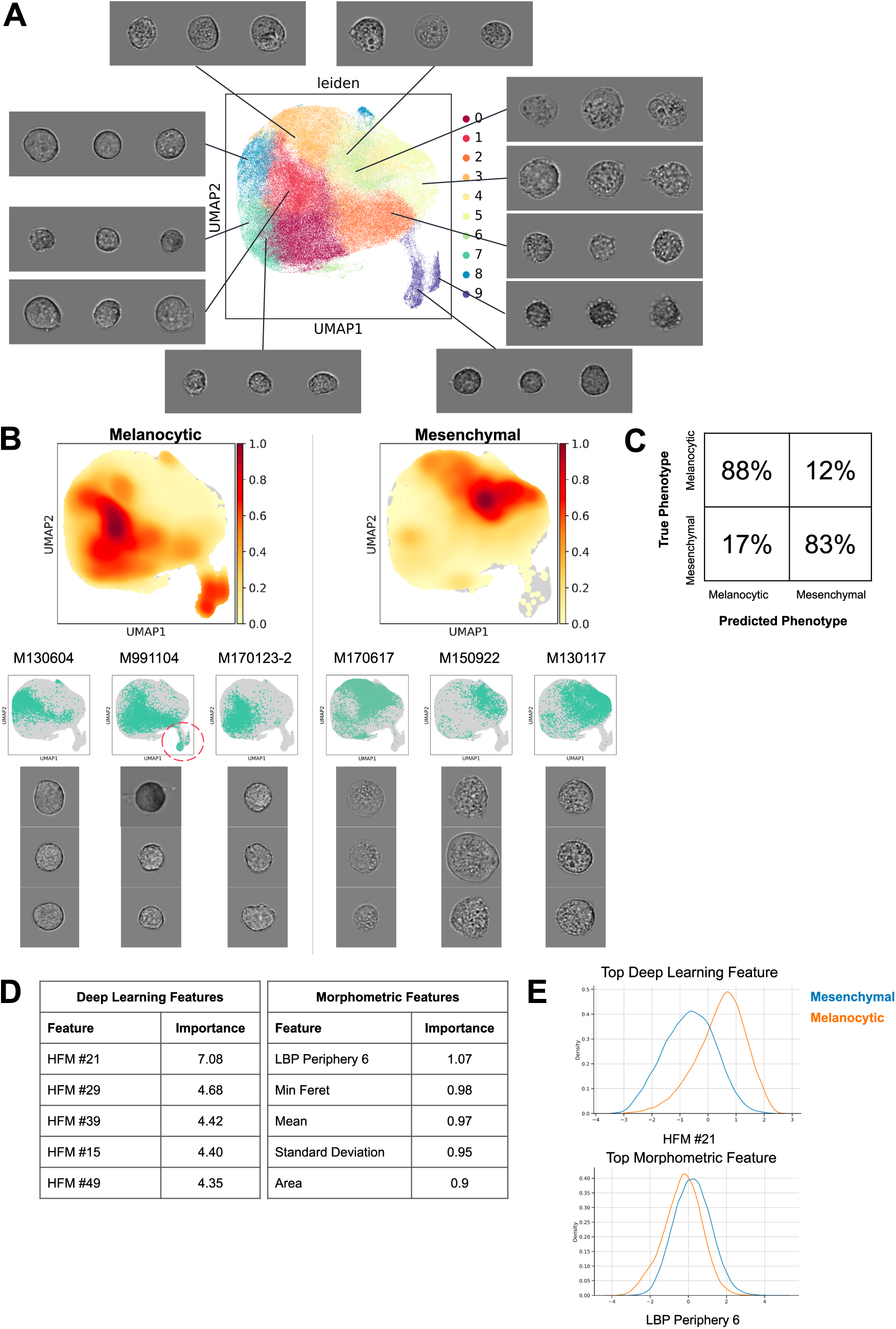
High-dimensional morphology distinguishes melanocytic and mesenchymal cell lines. **(A)** Morphology analysis was performed on 19 melanoma cell lines (listed in Table 1) and visualized in reduced-dimensional space as a UMAP. The UMAP is colored by Leiden clustering of the morphology features. Representative images from each Leiden cluster are shown, which demonstrated the morphological diversity in the melanoma cell lines, including a visually distinct pigmented population (Leiden Cluster 9). **(B)** The UMAPs were colored by relative density, and unique morphotypes for melanocytic and mesenchymal cells emerged. UMAPs showing the distribution of cells for individual samples from representative cell lines are shown, as well as representative images. UMAPs and images for all remaining cell lines can be found in Supplementary Figure 1. **(C)** The Melanoma Phenotype Classifier predicted the correct phenotype with up to 88% accuracy, with false negatives and false positives below 20%. **(D)** The top differential morphology features distinguishing melanocytic and mesenchymal cells are shown with the relative importance score. The top features were all deep learning features. The top morphometric features are shown as well, which represent texture (LBP Periphery), intensity (Mean, Standard Deviation) and size-based features (Min Feret, Area). Definitions of all features can be found in Supplementary Table 1. **(E)** The distributions of the top deep learning and morphometric features are shown for melanocytic and mesenchymal cell types.

In order to determine the top morphology features distinguishing the two phenotypes, we used a Random Forest Model (38) to reveal that the top features distinguishing the two phenotypes were mainly deep learning features (DL21, DL29, DL39, DL15, and DL49) (Fig. 2D), which are not readily human-interpretable. To provide interpretability to this analysis, we examined the top morphometric (human-interpretable) features distinguishing the two phenotypes (Fig. 2E, Supplementary Table 5), which revealed a mixture of texture and sized-based features: Local binary pattern (LBP) Periphery 6, Min Feret, Mean, Standard Deviation, and Area. These results were supported by visual inspection of representative cell images (Fig. 2B). In sum, our morphology analysis revealed that mesenchymal cells are larger and more granular on average relative to melanocytic cells.

### Classifier predictions applied to phenotypic shifts induced by perturbations

Melanoma cells exhibit the ability to shift from one phenotype to another upon chemical or genetic perturbations, which is a mechanism of therapeutic resistance and metastasis induction (27,39). To demonstrate the utility of the Melanoma Phenotype Classifier to detect induced phenotype shifts, we imaged control and perturbed cells on the Deepcell platform, and used the Melanoma Phenotype Classifier to quantify the phenotype shift induced by each perturbation. Classification results were confirmed by scRNA-Seq (Fig. 3A). The cell lines selected for this part of the study were not part of the classifier training set, which meant that the classifier had not previously “seen” these cells.

**Figure 3:**
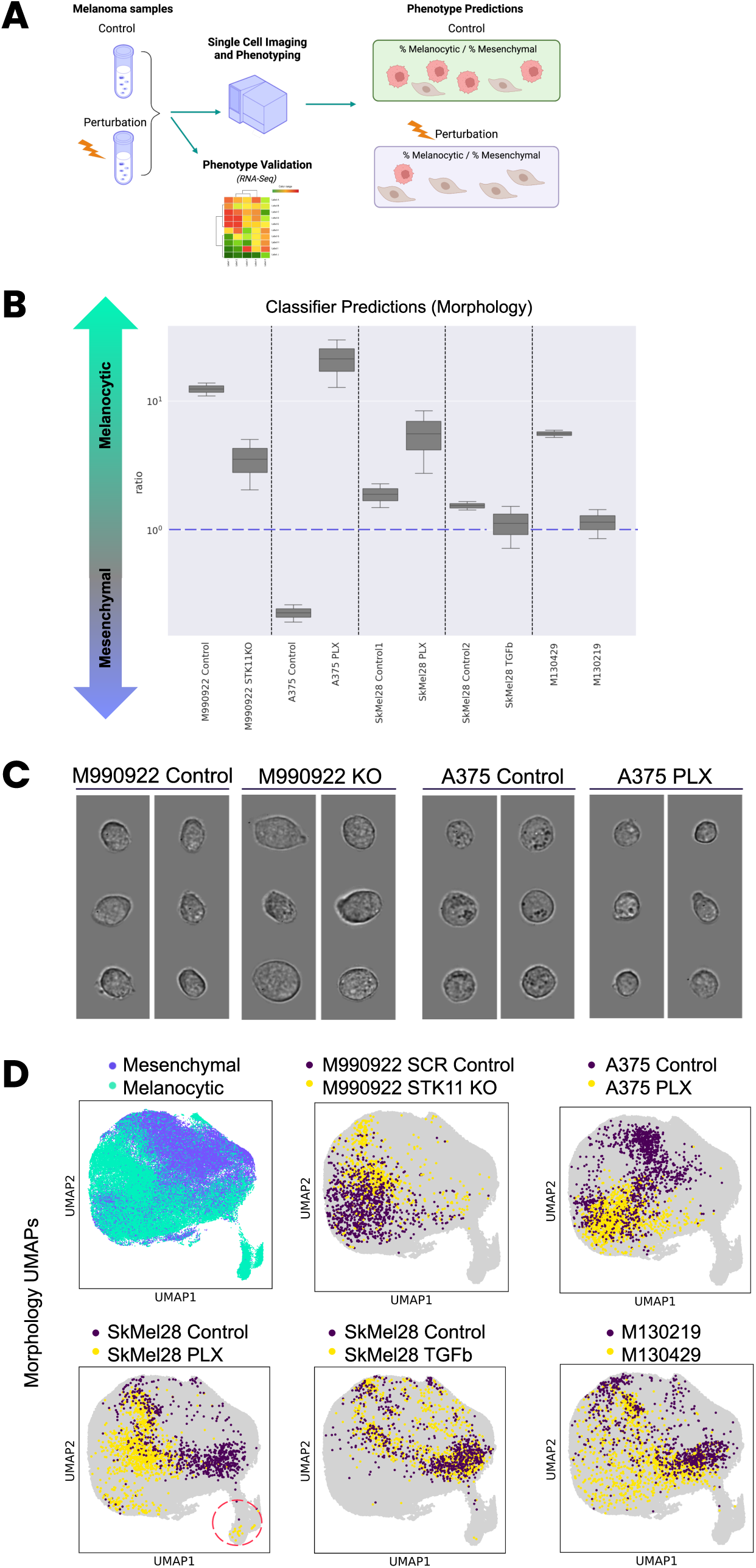
Morphometric analysis of phenotypic states induced by perturbations. **(A)** Control and perturbed cell lines were phenotyped by RNA-Seq and imaged on the Deepcell platform to generate phenotype predictions. **(B)** Measurement of the predicted melanocytic to predicted mesenchymal ratio for each control and perturbed cell line pair. Above 1, the cell populations are more melanocytic. Below 1, the cells are more mesenchymal. **(C)** Representative images of two pairs of control and perturbed cell lines, highlighting morphological changes induced by the perturbations that were used to predict phenotype. **(D)** UMAPs of morphology embeddings space for the five pairs of control and perturbed cell lines, enabling visualization of the morphology shifts induced by chemical and genetic perturbations. Detection of pigmented cells is indicated by the red circle. STK11KO = CRISPR-based knockout of STK11 gene, PLX = BRAF inhibitor (vemurafenib), TGFβ = transforming growth factor-beta.

To evaluate the induced shifts, we measured the ratio of the predicted melanocytic percentage to the predicted mesenchymal percentage (Fig. 3B). Ratios >10^0^ indicated a majority of melanocytic cells, while ratios <10^0^ indicated a majority of mesenchymal cells. For genetic perturbation, we used CRISPR to knock out the STK11 gene in M990922 cells. Knockout of this tumor suppressor gene has previously been shown to induce invasive properties (27), and morphological analysis by the Melanoma Phenotype Classifier revealed a distinct shift towards a mesenchymal state, represented by a 5.5x shift in the melanocytic : mesenchymal ratio. For chemical perturbation, we treated cells with an oncogenic BRAF inhibitor (PLX, vemurafenib) that has been shown to induce phenotypic shifts in various melanoma cell line models (14). In our study, treatment of A375 and SkMel28 cells with vemurafenib induced an expected shift toward a melanocytic phenotype, with an observed 77x ratio shift for A375 cells, and a 1.9x ratio shift for SkMel28 cells (Fig. 3B). A second perturbation was TGF-β, which induced a subtle, expected shift towards a mesenchymal state with a corresponding ratio shift of 1.5x. We next applied the classifier to two cell lines originating from the same patient, but exhibiting opposite phenotypes. Despite their identical genetic backgrounds, the Melanoma Phenotype Classifier correctly identified M130429 cells as melanocytic and M130219 cells as less melanocytic with a ratio shift of 4.3x (26). All of the predicted shifts generated by the classifier were confirmed by scRNA-Seq (Supplementary Fig. 3A-B) or by bulk RNA-Seq (M130219 and M130429 cell lines, Supplementary Fig. 4A). Morphological changes induced by the perturbations could be observed in the high resolution images (Fig. 3C, Supplementary Fig. 4). For example, the STK11 KO M990922 cells were larger on average than the corresponding M990922 control cells, while A375 cells treated with vemurafenib were smaller and exhibited less black granules on average than control A375 cells (Supplementary Fig. 4). To further interrogate phenotype shifts induced by each perturbation, we generated UMAPs of the embeddings of the control cells and corresponding perturbed cells (Fig. 3D, Supplementary Fig. 5). These plots corroborated the predicted shifts produced by the Melanoma Phenotype Classifier, and additionally revealed the presence of cell subpopulations induced by some of the perturbations. For example, pigmented cells were observed among SkMel28 cells treated with vemurafenib (Fig. 3D, red circle). Thus, the Melanoma Phenotype Classifier successfully predicted phenotype switching for a variety of cell lines and perturbations within the same cell lines, which was confirmed by scRNA-Seq gene signature analysis.

### Predicting phenotype variability in clinical samples with the classifier

To demonstrate its utility on clinical samples and its potential orthogonality to scRNA-Seq, we evaluated the Melanoma Phenotype Classifier on six dissociated tumor cell (DTC) samples (Supplementary Table 3). We plotted density UMAPs of the morphology embeddings of melanocytic and mesenchymal cell lines adjacent to UMAPs of scRNA-Seq data from melanocytic and mesenchymal cell lines based on transcriptional gene sets of each phenotype (Fig. 4A). Morphology and transcriptional UMAPs exhibited similar patterns in this morphology space, highlighting the uniqueness of the two phenotypes and the potential orthogonality of the two approaches. These UMAPs also served as references for the UMAPs generated for individual DTCs (Fig. 4B), revealing that most samples occupied similar areas of morphology space and transcriptional space. The representative cell images of each DTC revealed that these cells were on average significantly smaller than the 19 melanoma cell lines analyzed in this study (Fig. 2, Fig. 3). These images and the corresponding UMAPs also revealed that DTC 3 had a subpopulation of pigmented cells (Fig. 4B, red circle, red arrows).

**Figure 4:**
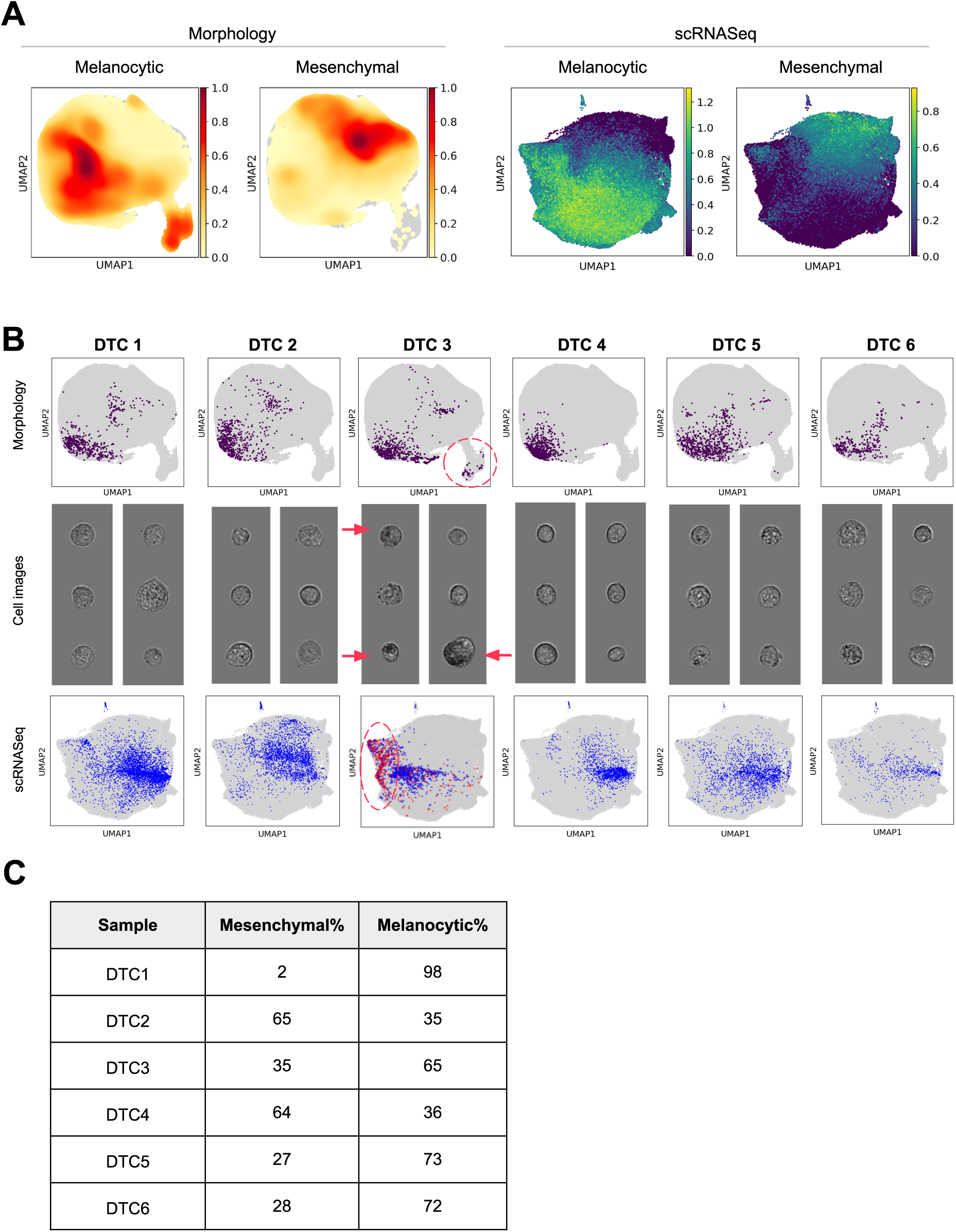
Morphology analysis, phenotype prediction, and scRNA-Seq analysis of dissociated tumor cell (DTC) samples. **(A)** Density UMAPs of morphology embeddings space (left) and transcriptional space based on scRNA-Seq (right) exhibit distinct localization of the two melanoma phenotypes based on cell line data. **(B)** UMAPs of morphology embedding space for each DTC (top), representative cell images from each DTC (middle), and UMAPs of scRNA-Seq for each DTC (bottom. Red circle indicates detection of pigmented cells in DTC3, which are identified by the red arrows in the representative cell images. **(C)** Percentages of the predicted melanocytic vs. mesenchymal phenotypes based on morphology for each DTC.

We next evaluated the phenotype predictions generated by the Melanoma Phenotype Classifier on the 6 DTCs and compared them to the phenotypes determined by melanocytic and mesenchymal gene sets from corresponding scRNA-Seq data. The prediction percentages for each DTC were compiled (Fig. 4C), along with the phenotype percentages determined by scRNA-Seq (Supplementary Table 7). The classifier correctly predicted that the samples were mostly melanocytic for 4 of the 6 DTCs (DTCs 1, 3, 5, and 6). For DTC 4, the classifier predicted that the sample was a mix of both phenotypes (64% Mesenchymal, 36% Melanocytic), which was also supported by the scRNA-Seq data (44% Mesenchymal, 56% Melanocytic). The main discrepancy between the classifier predictions and scRNA-Seq was observed with DTC 2, for which the classifier also predicted a mix of phenotypes (65% Mesenchymal, 35% Melanocytic), but scRNA-Seq indicated a majority of cells were melanocytic (1% Mesenchymal, 99% Melanocytic). Overall, these results demonstrated that there was good concordance between the phenotype predictions from the Melanoma Phenotype Classifier and the phenotypes determined by scRNA-Seq-based marker gene sets.

## Discussion

In this study, we applied Deepcell technology to develop a high-accuracy Melanoma Phenotype Classifier to enable label-free phenotyping at the single-cell level. We trained the classifier using cell lines with known melanoma phenotypes based on transcriptional signatures, achieving an accuracy exceeding 80% on cell lines that had not been used for training. Furthermore, we demonstrated the ability of the classifier to identify phenotypic shifts induced by chemical or genetic perturbations, which were validated by scRNA-seq. Importantly, we also illustrated its applicability in assessing the phenotypes of clinical samples, specifically dissociated tumor cell samples from melanoma patients. Our findings show that morphology accurately captures complex phenotype information. We foresee that morphotyping (phenotyping based on morphology) will enable label-free single-cell phenotype-based classification and ultimately sorting of live melanoma cells.

Phenotype switching of melanoma cells is prevalent during tumor evolution, and may lead to drug resistance. The use of precise phenotyping methods may facilitate the development of more precise personalized therapies in melanoma or other tumor types (4,40). The Melanoma Phenotype Classifier developed in this study offers a label-free approach to cancer phenotyping, utilizing morphology to distinguish between melanocytic and mesenchymal cell states. Although visual differences have been reported between different melanoma cell states (22,23), to our knowledge this is the first label-free classifier that predicts melanoma phenotypes based on visual features. Although we could not perform a direct comparison with analogous studies, the >80% accuracy achieved by the label-free classifier surpasses a previous machine learning approach that differentiated low (B16-F1) and high (B16F-10) metastatic murine melanoma cells with ∼74% accuracy using F-actin labeling (41). The accuracy of phenotype prediction within cell lines showed slight variation, especially in melanocytic cell lines, ranging from 60.4% to 97.1%. This variance in prediction accuracy for individual cell phenotypes could be attributed to the existence of cells in transitional or intermediate cell states, or to the fact that the cell lines were not derived from single-cell clones and may exhibit some heterogeneity in cell states. Phenotype switching is not a binary transition and both melanocytic and mesenchymal substates have been reported (42–45). Differences may also stem from limited concordance between phenotyping methods. In our study we relied on transcriptional profiling to define the ground truth. However, phenotypes are influenced by factors that may not be captured by RNA-seq, such as post-transcriptional or post-translational modifications, and environmental factors. Indeed, there are discrepancies between functional studies and transcriptional profiling. For example, conditional deletion of the inhibitory signaling factor Smad7 leads to melanoma cells acquiring both proliferative and invasive properties, thus uncoupling phenotype switching from metastatic behavior (46). Analogously, morphological studies and transcriptional profiling are not fully consistent (47).

We also demonstrated the feasibility of applying the Melanoma Phenotype Classifier to clinical samples, but with diminished accuracy in sample prediction (correctly classifying five out of six DTCs) compared to cell line samples. This could be attributed to a couple of factors. Firstly, the Melanoma Phenotype Classifier was exclusively trained with cell line samples, which exhibit less phenotypic and morphological variability under in vitro cell culture conditions. Secondly, melanoma cells were not preselected prior to the direct application of the classifier to DTCs, and the presence of fibroblasts and endothelial cells may have led to skewing towards a mesenchymal phenotype. To enhance applicability to clinical specimens, future iterations will involve training the classifier with clinical samples. Expansion of training sets to include cell lines of intermediate phenotypes (Supplementary Fig. 2), and clinical samples from a wider diversity of tumor sites and treatment regimens should improve robustness and applicability of the classifier. The novel use of the HFM as a framework for classifier development will enable rapid future updates to the Melanoma Phenotype Classifier. Considering the increased complexity and heterogeneity of melanoma phenotypes in vivo, refining our model to identify subtle subtypes may also have clinical implications, particularly in the context of treatment resistance, as indicated by recent research linking the mesenchymal phenotype with immune checkpoint inhibitors (ICI) resistance (17).

A key finding from analysis of our Morphology UMAPs was the ability to identify subpopulations of pigmented cells, which have been reported to have an impact on melanoma progression and drug response (48). These subpopulations appeared in control cell lines (Fig. 2B), perturbed cell lines (Fig. 3D), and in DTCs (Fig. 4B). The ability to analyze pigmented cells with label-free methods may be useful for studying the process of pigment production and associated diseases (49).

Remarkably, the Melanoma Phenotype Classifier was able to detect phenotypic changes resulting from a single CRISPR-induced mutation (Fig. 3B, 3D). This suggests the possibility of combining high-throughput CRISPR perturbation screens (e.g., perturb-Seq) with morphological profiling (50). Identifying correlations between specific mutations and distinct morphological phenotypes, combined with the ability to sort them, could significantly enhance our understanding of the functional implications of genetic alterations in melanoma.

In conclusion, the presented Melanoma Phenotype Classifier holds significant potential for a variety of applications, advancing both biological and clinical research in melanoma. Having demonstrated the performance of morphology-based phenotyping of melanoma cells, future directions of this research should include application of these methods to better elucidate associations between phenotype and clinically relevant features. The use of a label-free approach preserves cell viability, thus enabling downstream sorting of cells with the desired melanoma phenotype. For example, phenotype-based sorting of clinical melanoma samples may be combined with orthogonal approaches such as metabolomics, to assess metabolomic requirements of clinically relevant melanoma phenotypes. Combination of morphological characterization with more comprehensive genomics methods should be explored to identify associations between specific phenotypes and known clinically relevant biomarkers, which would be particularly interesting for patients who suffer from melanoma lacking BRAF mutations or are resistant to ICIs. Morphology-based phenotyping may also be applied more directly to assess drug resistance or metastatic potential of specific melanoma phenotypes. Within the broader context of oncology, our approach paves the way for morphotyping-based classification and sorting of a wide range of tumor cells beyond melanoma, and thus presents a cost-effective alternative to traditional transcriptomic methods.

## Acknowledgments

The authors would like to thank the URPP biobank at the University of Zurich for access to the melanoma cell lines, as well as Senzeyu Zhang, Ryan Carelli, Kevin Jacobs, Nianzhen Li, Mahyar Salek, and Maddison Masaeli for their contributions to the study. We also thank various Deepcell team members for hardware and software support, as well as reviewing the manuscript prior to submission. Figure 1 was created with BioRender.

