## Supplementary Tables for "Label-Free Melanoma Phenotype Classification Using Artificial Intelligence-Based Morphological Profiling"

**Supplementary Table 1: List of Commercial Cell Lines**

| **Cell Line** | **Vendor/Source** | **Catalog No.** | **Tumor Site** | **Driver Mutation** | **Age (years)** | **Sex (Female: F; Male: M)** | **Usage** |
| --- | --- | --- | --- | --- | --- | --- | --- |
| SkMel28 | ATCC | HTB-72 | Skin | BRAF (V600E) | 51 | M | Classification |
| A375 | ATCC | CRL-1619 | Skin | BRAF (V600E) | 54 | F | Classification |
| MNT-1 | ATCC | CRL-3450 | Lymph node | BRAF (V600E) | N/A | N/A | Training/Validation |
| WM983A | Wistar Institute | CVCL_6808 | Primary site | BRAF (V600E) | 54 | M | Training/Validation |

**Supplementary Table 2: List of Primary Cell Lines** all primary cell lines were acquired from the URPP biobank at the University of Zurich from consenting patients of Caucasian origin and age between 29 and 84 years; 15 male patients and 13 female patients.

| **Cell Line** | **Tumor Site** | **Driver Mutation** | **Age (years)** | **Sex (Female: F; Male: M)** | **Usage** |
| --- | --- | --- | --- | --- | --- |
| M130219 | Subcutaneous | NRAS(Q61R) | 48 | M | Classification |
| M130820 | Lymph node | BRAF (V600E) | 65 | M | Classification |
| M150818 | Lymph node | BRAF (V600E) | 46 | F | Classification |
| M140307 | Lymph node | BRAF (V600K), NRAS (G12A) | 74 | M | Training/Validation |
| M120815 | Lymph node | NRAS (Q61R) | 84 | F | Training/Validation |
| M161201 | Lymph node | BRAF (V600E) | 67 | M | Training/Validation |
| M170123_2 | Lymph node | WT | 71 | F | Training/Validation |
| M160411 | Lymph node | BRAF (V600-K601delinsE) | 45 | F | Training/Validation |
| M980513 | Lymph node | BRAF (V600E) | 32 | F | Training/Validation |
| M991104 | Lymph node | BRAF (V600E) | 30 | F | Training/Validation |
| M130604 | Lymph node | BRAF (V600K) | 72 | M | Training/Validation |
| M180212 | Lymph node | BRAF (V600E) | 45 | F | Classification |
| M150922 | Lymph node | BRAF (V600E) | 63 | M | Training/Validation |
| M990922 | Lymph node | BRAF (V600E) | 29 | F | Classification |
| M170617 | Colon | NRAS (Q61R) | 64 | F | Training/Validation |
| M150536 | Brain | BRAF (V600E) | 65 | M | Training/Validation |
| M130429 | Bone | NRAS(Q61R) | 48 | M | Classification |
| M130515 | Lymph node | NRAS (Q61R) | 58 | M | Classification |
| M111031 | Lymph node | BRAF (V600E) | 33 | M | Classification |
| M121224 | Thorax | BRAF (V600E), NRAS (Q61K) | 37 | M | Classification |
| M130903 | Lymph node | BRAF (V600E), NRAS (Q61H) | 57 | M | Classification |
| M150423 | Liver | BRAF (V600E), NRAS (Q61R) | 56 | M | Classification |
| M130117 | Pectoral/axial left | NRAS (Q61R) | 48 | M | Training/Validation |
| M150672 | Thorax | BRAF (V600E) | 41 | F | Training/Validation |
| M130107 | Back | NRAS (Q61L) | 65 | F | Training/Validation |
| M170314 | Lymph node | BRAF (V600E) | 66 | F | Training/Validation |
| M150548 | Brain | BRAF (V600E) | 65 | M | Training/Validation |
| M160526 | Lymph node | BRAF (V600-K601delinsE) | 45 | F | Training/Validation |

**Supplementary Table 3: List of dissociated tumor cell samples (DTCs)** all patient biopsies were purchased from Discovery Life Sciences: all patient biopsies were of Caucasian origin, with ages ranging from 42 to 87 years, and 4 female patients and 2 male patients.

| **Name** | **Sample ID** | **Tumor site** | **Clinical Stage** | **Age (years)** | **Sex (Female: F; Male: M)** | **Usage** |
| --- | --- | --- | --- | --- | --- | --- |
| DTC 1 | BTC1000-5F130512285101220MS | Lymph Node | III-D | 61 | F | Classification |
| DTC 2 | BTC1000-7G200015995081621MS | Unknown | Unknown | 43 | F | Classification |
| DTC 3 | BTC1000-G9121494809102119MS | Unknown | Unknown | 54 | M | Classification |
| DTC 4 | BTC1000-5F200020387110221MS | Skin | III-C | 87 | M | Classification |
| DTC 5 | BTC1000-J7110002996020317MS | Lymph Node | III | 57 | F | Classification |
| DTC 6 | BTC1000-5F200009946072020MS | Lymph Node | III-B | 42 | F | Classification |

**Supplementary Table 4: Phenotype Gene List**

| **Mesenchymal** | **Melanocytic** |
| --- | --- |
| *AQP1* | *BIRC7* |
| *AXL* | *CAPN3* |
| *BGN* | *CCL2* |
| *CDH13* | *CCL3* |
| *CDH2* | *CCL4* |
| *COL5A1* | *CCL5* |
| *CTGF* | *CCL8* |
| *CYR6* | *CCL9* |
| *EGFR* | *CDH19* |
| *FBN1* | *CSPG4* |
| *FEZ1* | *DCT* |
| *FGF2* | *ERBB3* |
| *FOSL2* | *FXYD3* |
| *GFRA2* | *GJB1* |
| *GFRA3* | *GPM6B* |
| *IGFBP6* | *GPR143* |
| *IL6* | *IL23* |
| *INHBA* | *LINC00473* |
| *L1CAM* | *LOXL4* |
| *LOX* | *MAGEA6* |
| *LOXL2* | *MELTF* |
| *NGFR* | *MIA* |
| *PDGFRB* | *MLANA* |
| *POSTN* | *MLPH* |
| *RGS5* | *PAX3* |
| *RSPO3* | *PLEKHB1* |
| *SERPINE1* | *PLP1* |
| *SLC22A17* | *PMEL* |
| *SLC2A1* | *PRAME* |
| *SLIT2* | *QPCT* |
| *SLITRK6* | *RAB38* |
| *SNRNP25* | *S100A1* |
| *SOX9* | *S100B* |
| *SPIB* | *SEMA3B* |
| *SUSD1* | *SLC24A5* |
| *TAGLN* | *SLC26A2* |
| *TCF4* | *SLC45A2* |
| *TCL1A* | *SOX10* |
| *TGFBI* | *TRPM1* |
| *TGM2* | *TYR* |
| *THBS1* | *TYRP1* |
| *TMEM176B* |  |
| *TNC* |  |
| *TNFRSF11B* |  |
| *TPM1* |  |
| *TPM2* |  |
| *TSPAN13* |  |
| *UGCG* |  |
| *VIM* |  |
| *WNT5A* |  |

**Supplementary Table 5: Definitions of Morphometric Features**

| **Category** | **Morphometric** | **Definition** |
| --- | --- | --- |
| Geometry | AREA | Cell area (µm²). |
| Geometry | PERIMETER | Length of the cell outline (µm). |
| Geometry | MAX_FERET | Length of longest Feret diameter (longest caliper measurement) (µm). |
| Geometry | MIN_FERET | Length of shortest Feret diameter (shortest caliper measurement) (µm). |
| Geometry | MAX_RADIUS | Largest radius (µm). |
| Geometry | MIN_RADIUS | Smallest radius (µm). |
| Geometry | LONG_AXIS | Long axis of best fit ellipse (µm). |
| Geometry | SHORT_AXIS | Short axis of best fit ellipse (µm). |
| Geometry | ECCENTRICITY | Aspect ratio of best fit ellipse (unitless). |
| Geometry | ELLIPSE_VARIANCE | Ellipse variance, deviation from elliptical shape (unitless). |
| Geometry | ROUNDNESS | Similarity to a circle (unitless). |
| Geometry | CIRCULARITY | Coefficient of variation of the radius (unitless). |
| Geometry | SOLIDITY | Ratio of the area of the convex hull of the cell to the area of the cell (unitless). |
| Intensity | MEAN | Mean pixel grayscale value. |
| Intensity | STANDARD_DEVIATION | Standard deviation of pixel grayscale values. |
| Intensity | POSITIVE_FRACTION | Fraction of pixels with a gray value above 0.1. |
| Intensity | NEGATIVE_FRACTION | Fraction of pixels with a gray value below -0.1. |
| Intensity | PERCENTILE_75 | 75th percentile of pixel grayscale values. |
| Intensity | PERCENTILE_25 | 25th percentile of pixel grayscale values. |
| Intensity | SMALL_WHITE_BLOBS_COUNT | Number of small white blobs. |
| Intensity | SMALL_BLACK_BLOBS_COUNT | Number of small black blobs. |
| Intensity | LARGE_WHITE_BLOBS_COUNT | Number of large white blobs. |
| Intensity | LARGE_BLACK_BLOBS_COUNT | Number of large black blobs. |
| Intensity | SMALL_WHITE_BLOB_INTEGRAL | Integral over small white blobs. |
| Intensity | SMALL_BLACK_BLOB_INTEGRAL | Integral over small black blobs. |
| Intensity | LARGE_WHITE_BLOB_INTEGRAL | Integral over large white blobs. |
| Intensity | LARGE_BLACK_BLOB_INTEGRAL | Integral over large black blobs. |
| Texture | LBP_CENTER_{n} | Array of 10 uniform rotation-invariant Local Binary Pattern features, computed in the central region of the cell. |
| Texture | LBP_PERIPHERY_{n} | Array of 10 uniform rotation-invariant Local Binary Pattern features, computed in the peripheral region of the cell. Points = 8, radius = dc_cellimage.constants.PERIPHERY_LBP_RADIUS. The width of the periphery is determined through dc_cellimage.constants.PERIPHERY_WIDTH. |
| Moments | HU{n} | Array of 2 rotation and scale invariant weighted averages (moments) of the image pixel intensities. |

**Supplementary Table 6: scRNASeq-based Classification of cell lines**Measurement of the scRNASeq-derived values for the melanocytic and mesenchymal percentages for a subset of cell lines from Table 1.

| **Cell Line** | **Melanocytic%** | **Mesenchymal%** |
| --- | --- | --- |
| M140307 | 100% | 0 |
| M150548 | 100% | 0 |
| M170123_2 | 99% | 0 |
| M120815 | 1% | 88% |
| M130107 | 0 | 98% |
| M150922 | 0 | 99% |
| M170314 | 0 | 100% |

**Supplementary Table 7:** **Measurement of the scRNASeq-derived values for the melanocytic and mesenchymal percentages for each DTC**

| **Sample** | **Mesenchymal%** | **Melanocytic%** |
| --- | --- | --- |
| DTC1 | 1 | 99 |
| DTC2 | 1 | 99 |
| DTC3 | 1 | 99 |
| DTC4 | 44 | 56 |
| DTC5 | 3 | 97 |
| DTC6 | 1 | 99 |
