## Supplementary Figures for "Label-Free Melanoma Phenotype Classification Using Artificial Intelligence-Based Morphological Profiling"

Lattmann et al Supplementary Information

Supplementary Figures S1-S5

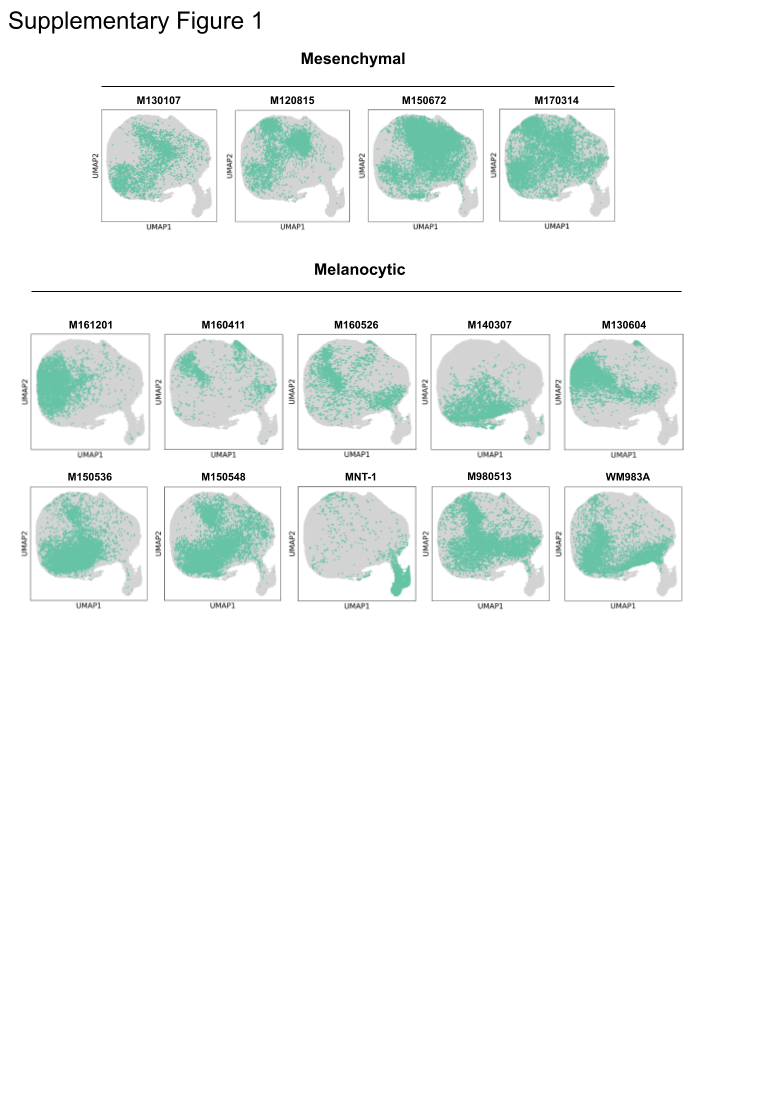

**Supplementary Figure S1: Individual UMAP plots of other mesenchymal and melanocytic cell lines used for classifier training.** Associated with Figure 2B.

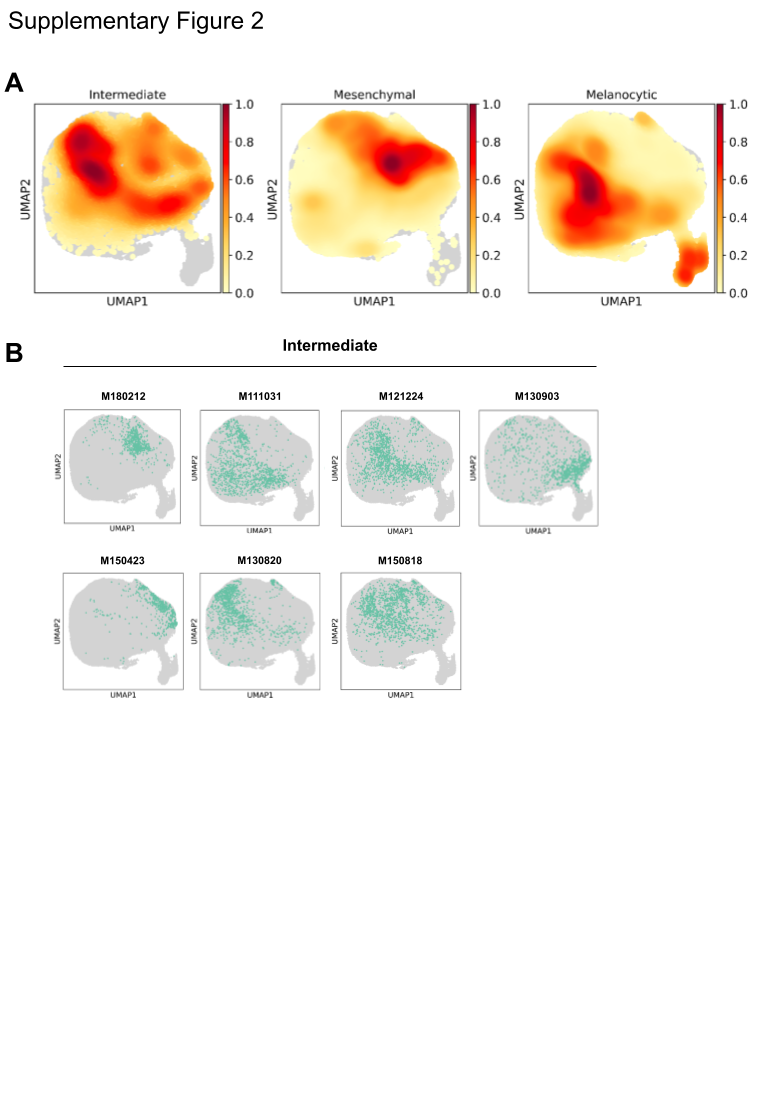

**Supplementary Figure S2: Phenotype density representation of melanoma cells with intermediate, mesenchymal and melanocytic phenotypes (A)** Aggregated density UMAP plots of intermediate, mesenchymal, and melanocytic cell lines. **(B)** Individual UMAP plots of all cell lines classified as “intermediate” with morphological signatures between mesenchymal and melanocytic. Associated with Figure 2B.

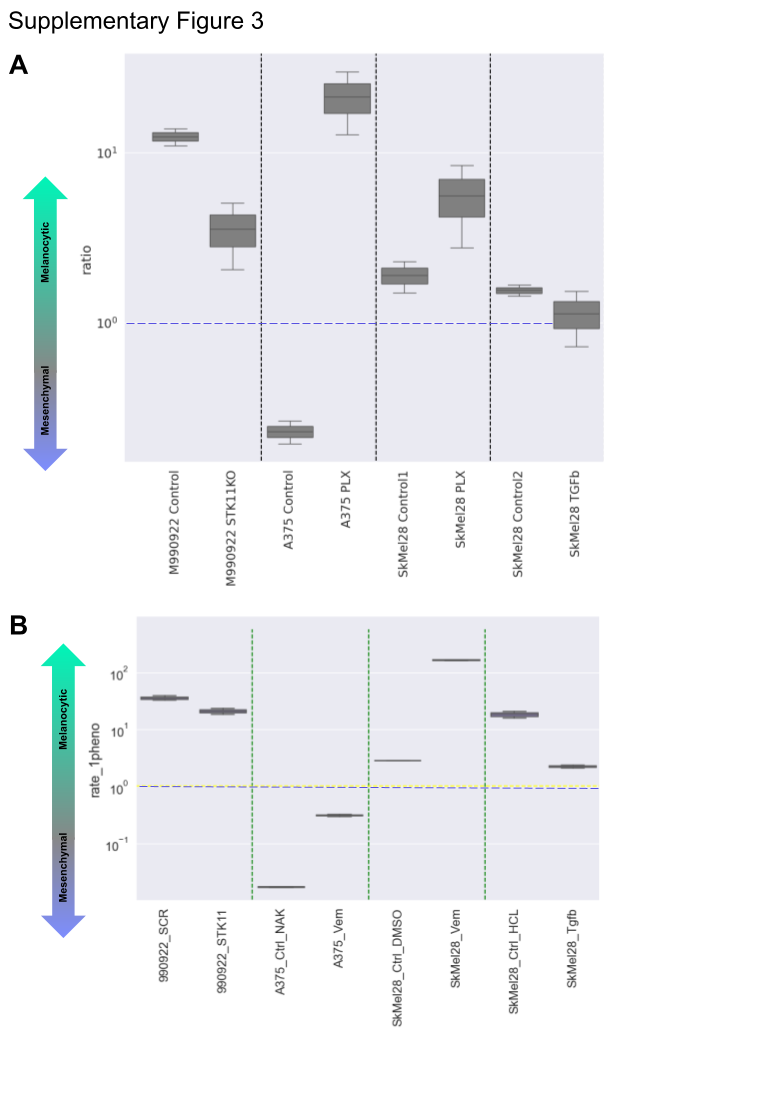
**Supplementary Figure S3: Confirmation of predicted phenotypic shifts by scRNA-Seq. (A)** Measurement of the predicted melanocytic to predicted mesenchymal ratio for each control and perturbed cell line pair. Above 1, the cell populations are more melanocytic. Below 1, the cells are more mesenchymal (same as Fig. 3B). **(B)** Measurement of the scRNASeq-derived values for the melanocytic to mesenchymal ratio for each control and perturbed cell line pair. For values above 1, the cell populations are more melanocytic. Below 1, the cells are more mesenchymal.

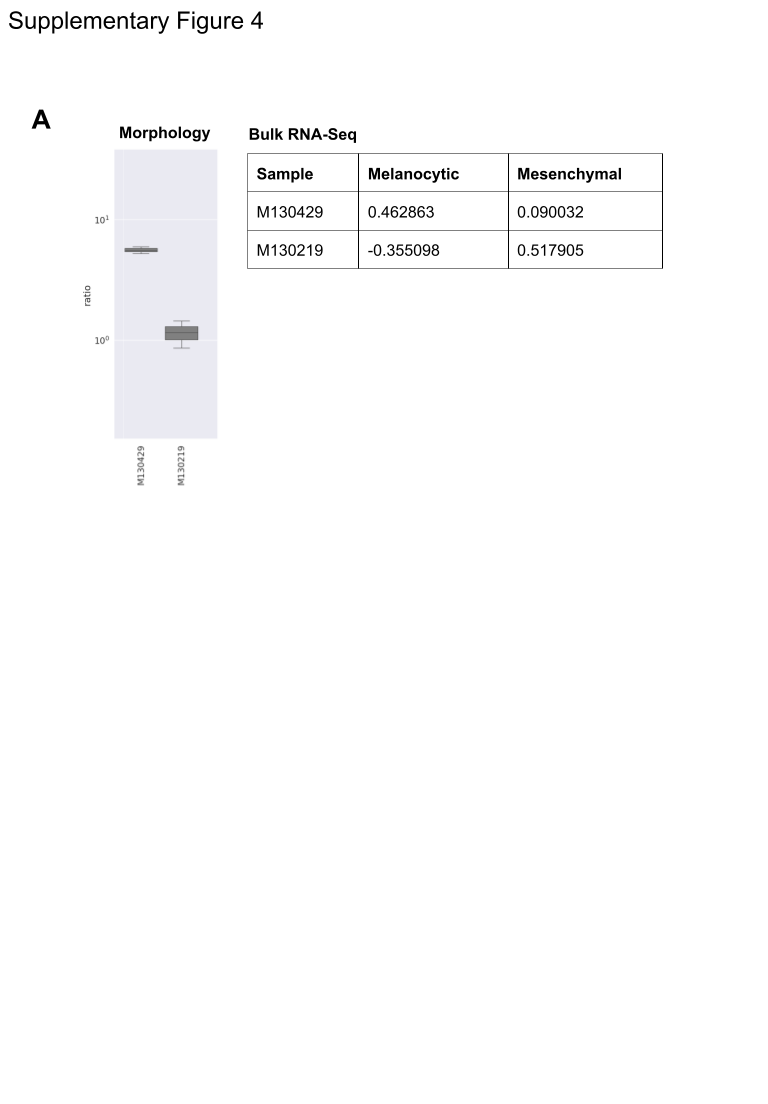

**Supplementary Figure S4: Bulk RNA-seq to confirm phenotypes of M130429 and M130219 cells derived from the same patient (A)** Measurement of the melanocytic to predicted mesenchymal ratio for the cell line pair M130429 and M130219 (left). Correlation coefficients against the melanocytic and mesenchymal phenotypes for each cell line based on bulk RNASeq data (right), using the genes in Supplementary Table 4.

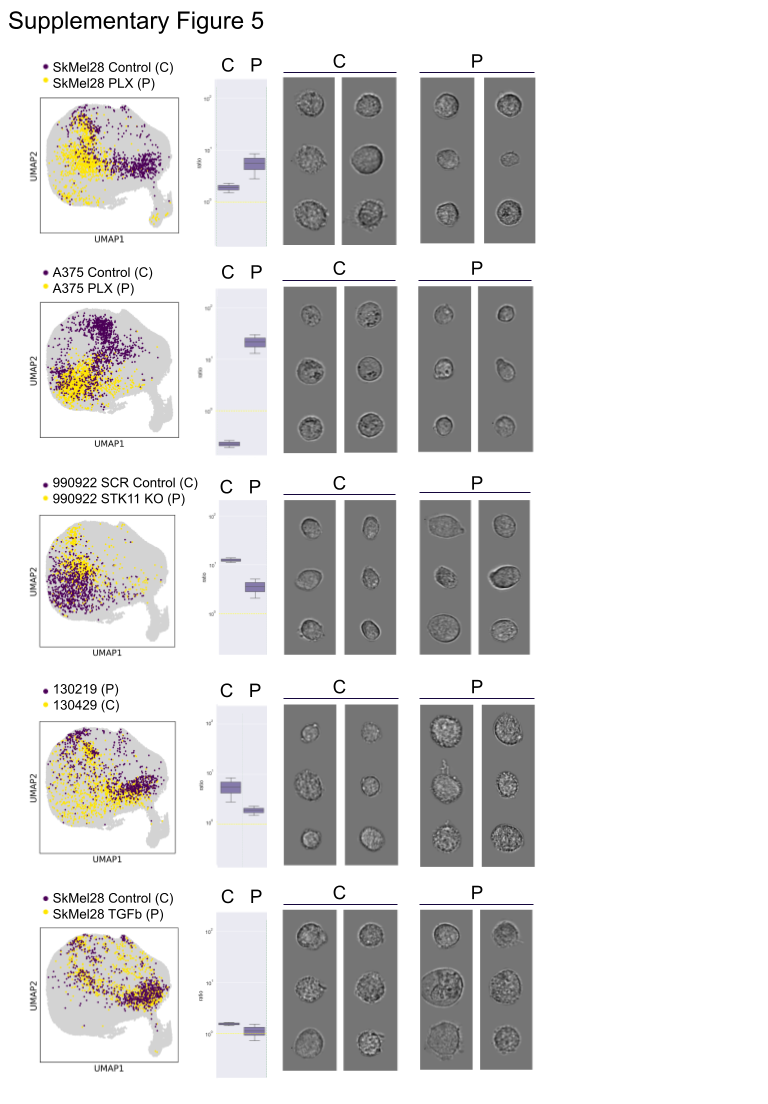

**Supplementary Figure S5: UMAPs of morphology embedding space for the five pairs of control and perturbed cell lines.** Enables visualization of the morphology shifts induced by chemical and genetic perturbations (left). Ratios of the predicted melanocytic to mesenchymal ratio for the control (C) versus perturbed (P) cells for each cell line pair (middle). Representative images of the control (C) and perturbed cells (P) for each cell line pair (right).
